# Extrusions promote engulfment and *Chlamydia* survival within macrophages

**DOI:** 10.1101/041079

**Authors:** Meghan Zuck, Tisha Ellis, Anthony Venida, Kevin Hybiske

## Abstract

All obligate intracellular pathogens must exit their host cells in order to propagate and survive as a species; the precise strategies they use have a direct impact on their ability to disseminate within a host, transmit to new hosts, and engage or avoid immune responses. The obligate intracellular bacterium *Chlamydia trachomatis* exits the host cell by two distinct exit strategies, lysis and extrusion. Despite being equally active pathways, lysis and extrusion differ greatly in their mechanisms. The defining characteristics of extrusions, and advantages gained by *Chlamydia* within this unique double-membrane structure are not well understood. Here, we present data that defines extrusions as being largely devoid of host organelles, comprised mostly of *Chlamydia* elementary bodies, and containing phosphatidylserine on the outer surface of the extrusion membrane. Towards defining a functional role for extrusions in *Chlamydia* pathogenesis, we demonstrate that extrusions confer significant infectious advantages for *Chlamydia* by serving as transient, intracellular-like niches for extracellular *Chlamydia*, as compared to *Chlamydia* that would otherwise exit by lysing the host cell. In addition to enhanced survival outside of the host cell, we report the key discovery that chlamydial extrusions can be engulfed by primary bone marrow-derived macrophages, after which they provide a protective microenvironment for *Chlamydia*. Extrusion-derived *Chlamydia* were able to stave off macrophage based killing beyond 8 h, and culminated in the release of infectious EB from the macrophage. Based on these findings, we propose a model in which a major outcome of *Chlamydia* exiting epithelial cells inside extrusions is to hijack macrophages as vehicles for dissemination within the host.

## Introduction

Chlamydiae are gram-negative obligate intracellular bacteria responsible for different diseases of clinical and public health importance. Among these are the causative agents of trachoma, the leading cause of infectious blindness worldwide [1], and the most prevalent bacterial sexually transmitted disease in the United States [2]. Infection is predominantly asymptomatic, and disease sequelae results from long-term infections or re-infections that can induce tissue damage and scarring [3]. In the absence of diagnosis and treatment, *Chlamydia trachomatis* infection can have severe outcomes, including infertility, ectopic pregnancy, and pelvic inflammatory disease [4,5]. Little mechanistic understanding exists for how *C. trachomatis* disseminate within a host, or transmit to new hosts, yet these mechanisms are likely complex given *Chlamydia*’s intracellular confinement and lack of inherent motility.

*Chlamydia* are characterized by a unique biphasic developmental cycle. Infection within a host begins with direct contact of *Chlamydia* elementary bodies (EB), the infectious and metabolically inert form of the bacteria, with columnar epithelial cells of mucosal tissue. Once the infectious EB has entered the cell, intracellular *Chlamydia* occupy a parasitophorous vacuole known as the inclusion. Within this protected intracellular niche, *Chlamydia* EB convert into the larger, metabolically active reticulate body (RB) and undergo successive rounds of replication until they asynchronously convert back into EB 24-48 h post infection, and ultimately exit the cell [4,6].

The ability to exit the host cell upon completion of intracellular developmental growth is a critical step in bacterial pathogenesis [7], and *Chlamydia* has evolved two mutually exclusive mechanisms for this task—lysis and extrusion [8]. Lysis is a destructive process, characterized by a sequential rupture of vacuole, nuclear and plasma membranes culminating in the release of free infectious bacteria [9]. Extrusion is a markedly different molecular mechanism of cellular exit, utilizing a packaged release strategy of *Chlamydia* that is defined by unique interactions between the bacteria and host cell [8]. Extrusion begins with invagination of the *Chlamydia*-containing inclusion, followed by a furrowing of the host cell plasma membrane to allow the pinching of the cell, resulting in the release of a membrane-encased compartment containing the *Chlamydia*, inclusion membrane, host cell cytoplasm, a meshwork actin network, and plasma membrane [8,10]. This process leaves the original host cell intact and often with a residual chlamydial inclusion. The lack of meaningful protective immunity in humans, and also the failure of efforts to generate efficacious vaccines despite the susceptibility of *Chlamydia* EB to antibody neutralization, highlight an innate biological capacity of *C. trachomatis* to suppress or elude the host immune system. The conservation of extrusion among chlamydiae is strongly suggestive of important roles in pathogenesis, such as dissemination within a host, transmission to new hosts, and may contribute to chronic infection through immune evasion [8].

Extrusions are novel pathogenic structures, and although there exists an understanding of the molecular pathways that govern the extrusion process [8,10-12], very little is known about their structure, internal composition, or functional roles in infection. In this study, we developed a method for enriching extrusions from infected cell cultures, which in turn enabled us to perform single cell characterization of the internal composition of these pathogenic structures. We found that extrusions are formed without the accompaniment of any host organelles that are otherwise associated with the infected host cell. Importantly, *C. trachomatis* derived from extrusions demonstrated enhanced extracellular survival compared to *C. trachomatis* released from host cells by lysis. Furthermore, we describe a novel process wherein *C. trachomatis* extrusions promote their engulfment, and subsequent survival, within primary bone marrow macrophages to eventually escape the macrophage while still retaining infectivity.

## Results

### Isolation and enrichment of *Chlamydia* extrusions

The current understanding of the composition of *Chlamydia* extrusions is they contain infectious bacteria that are enveloped by inclusion membrane, host cytosol, actin networks, and host plasma membrane [8,10]. In order to better characterize extrusions, potential functional benefits for extrusion-derived *Chlamydia*, and ultimately define the molecular mechanisms responsible for extrusion formation, a method for enrichment and separation of extrusions from host cells and extracellular *Chlamydia* is necessary.

We investigated a series of physical and pharmacological manipulations, and differential centrifugation strategies, to empirically derive a reliable method for the isolation and enrichment of extrusions from infected monolayers. We desired a process that was adaptable to other *C. trachomatis* strains and *Chlamydia* species, was independent of host cell type, and contained minimal handling. These efforts ultimately led to an approach in which endogenously produced extrusions were collected from supernatant of host cell monolayers, followed by sequential centrifugation steps, that resulted in an enriched suspension of extrusions, with reduced extracellular *Chlamydia*, host cells, and cellular debris (Fig 1A-B). Extrusions were routinely identified as circular, low contrast cellular objects, that lacked host nuclei and contained numerous bacteria. Although we cannot dismiss the possibility that some extrusions contain nuclei (e.g., when originating from multinucleated HeLa cells), this occurrence was rare and we chose to operationally define extrusions as *Chlamydia*-containing objects that lacked nuclei, in order to consistently differentiate them from infected host cells. There was visible heterogeneity in the appearance of extrusions by light microscopy, and some extrusions with high granularity were also typically present (Fig 1A).

**Fig 1.**
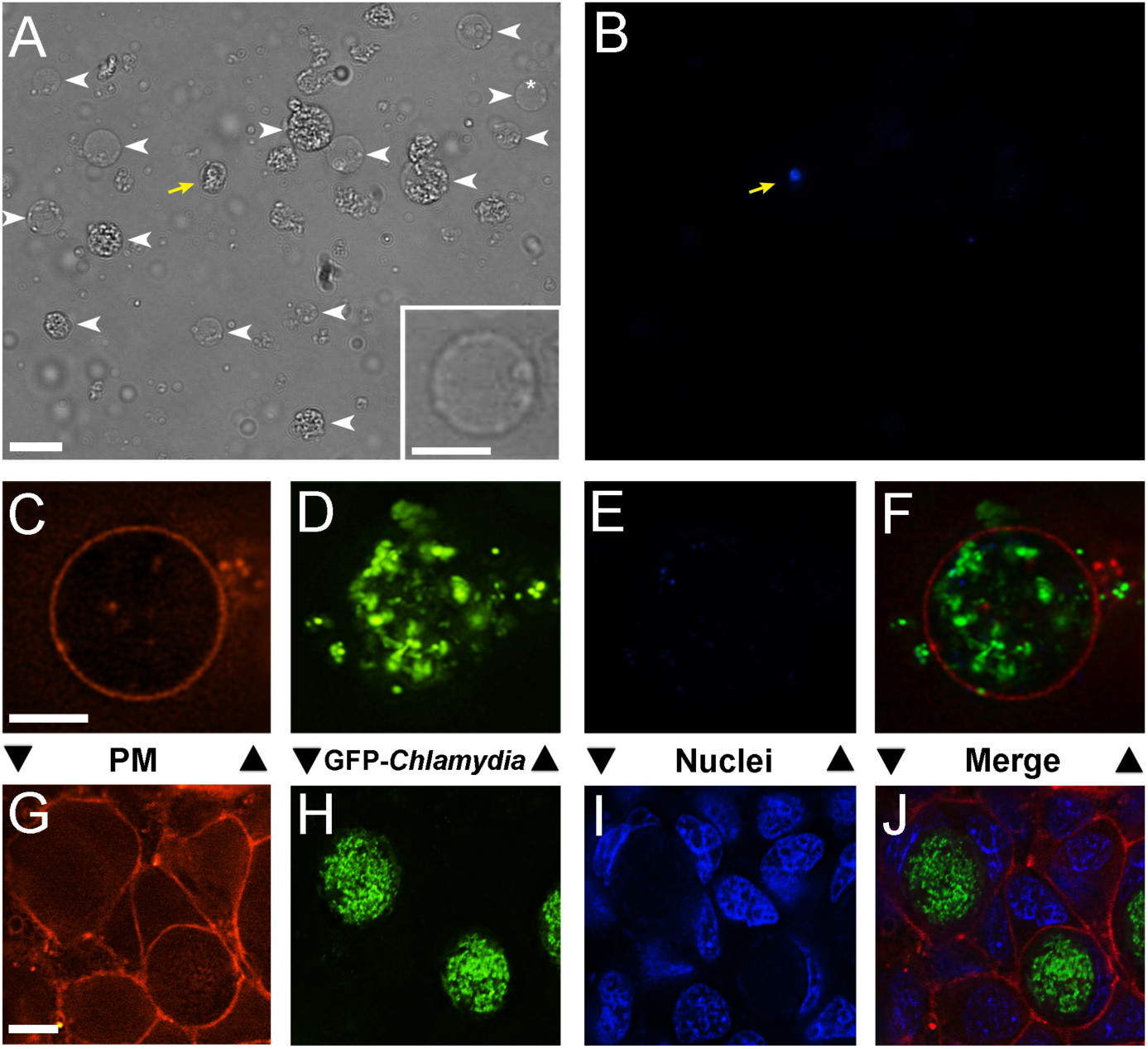
Isolation and enrichment of *Chlamydia* extrusions. **Top row:** representative field of extrusions following enrichment procedures, at 20× magnification. **A** Brightfield image showing 14 extrusions (white arrowheads). Inset: enlargement of the extrusion marked with white asterisk. **B** Micrograph of the same field of extrusions shown in **A**, labeled with DAPI (blue) to show the lack of nuclei in extrusions. A single host cell with nucleus is shown (yellow arrow). **Middle row:** representative micrographs of an isolated extrusion, visualized live with fluorescent probes for: **C** host plasma membrane (PM) (FM4-64, red), **D** GFP-expressing *C. trachomatis* (GFP, green), **E** nuclei (DAPI, blue), **F** merge of **C**-**E. Bottom row:** for comparison, micrographs of HeLa cells infected with GFP-expressing *C. trachomatis* L2 for 48 hpi, and visualized live with fluorescent probes for: **G** plasma membrane (FM4-64, red), **H** GFP-expressing *C. trachomatis* (green), **I** nuclei (DAPI, blue), **J** merge of **G**-**I**. Scale bar, 10 μm.

### Cellular composition of *Chlamydia* extrusions

We next sought to determine the internal and surface composition of *Chlamydia* extrusions. Because the mechanism of extrusion consists of contraction of the chlamydial inclusion and infected host cell [8], it was unknown whether extrusions additionally contained host organelles and other cellular structures. Alternatively, it was possible that the *Chlamydia-*directed extrusion mechanism might specifically result in a pathogenic compartment that contained only the chlamydial inclusion. We probed the composition of live extrusions by labeling them with specific fluorescent markers for key host structures, followed by live fluorescence microscopy. McCoy cells were infected with a *C. trachomatis* LGV serovar L2 strain transformed with a GFP expression plasmid [13], and extrusions were collected in a 2-4 h collection window at 48 hours post-infection (hpi). Enriched extrusions were typically circumscribed by host plasma membrane (Fig 1C), contained GFP-expressing *Chlamydia* (Fig 1D), and without host nuclei (Fig 1E). These data are in contrast to well-defined cell morphology of *C. trachomatis*-infected monolayers at 48 hpi (Fig 1G-J).

*Chlamydia* extrusions are morphologically similar to extracellular merosomes that are produced by malaria-infected hepatocytes, and that contain thousands of *Plasmodium* sporozoites [14-16]. The external surface of merosome membranes were shown to lack phosphatidylserine (PS), thus providing a potential mechanism by which extracellular malaria evade clearance by macrophages [15-17]. Given *Chlamydia*’s similar need to avoid immune clearance, we investigated whether PS was absent on *Chlamydia* extrusions, by labeling extrusions with fluorophore-conjugated annexin-V, which specifically binds to PS, and imaged a three dimensional *z*-series of extrusions by fluorescence microscopy. In contrast to *Plasmodium* merosomes, PS was frequently present on the surface of extrusions, typically as discrete puncta on the extrusion outer membrane (Fig 2A-D). A punctate PS pattern was quantitatively determined to be present on 59.6% of extrusions (Fig Q).

**Fig 2.**
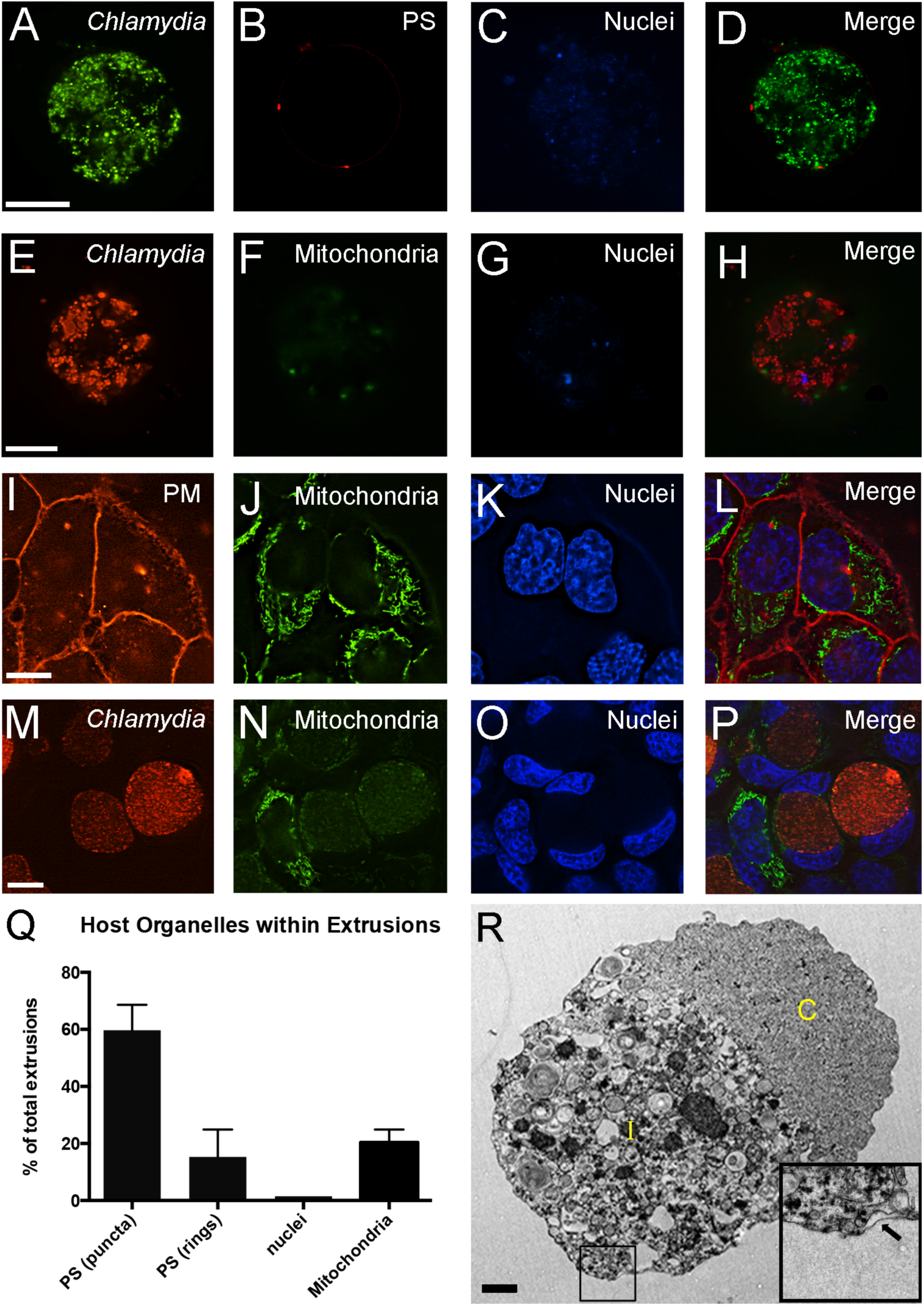
Host cell organelles are largely excluded from *Chlamydia* extrusions. **Top row:** representative micrographs of an isolated extrusion, visualized live with fluorescent probes for: **A** *C. trachomatis* (GFP, green), **B** phosphatidylserine (PS; annexin V-568, red), **C** nuclei (DAPI, blue), **D** merge of **A-C. Second row:** representative live extrusions showing: **E** *C. trachomatis* (mKate2, red), **F** mitochondria (mitotracker-488, green), **G** nuclei (DAPI, blue), **H** merge of **E-G. Third row:** micrographs of uninfected HeLa cells visualized live with probes for: **I** plasma membrane (PM) (FM4-64, red), **J** mitochondria (mitotracker-488, green), **K** nuclei (DAPI, blue), **L** merge of **I-K. Fourth row:** representative micrographs of HeLa cells infected with mKate2-expressing *C. trachomatis* for 48 h, and visualized live for: **M** *C. trachomatis* (mKate2, red), **N** mitochondria (mitotracker-488, green), **O** nuclei (DAPI, blue), **P** merge of **M-O. Q** Positive association of mitochondria, PS, and nuclei, was quantitatively enumerated from over 75 images of extrusions. Data points show mean + SEM, n = 3. **R** Transmission electron micrograph of a representative, isolated *C. trachomatis* extrusion at 0 hpe (hours post extrusion). Scale bar, 10 μm for **A-P**; 1 μm for **R**. Inset within **R** of enlarged region of extrusion showing presence of a double membrane on the periphery of the extrusion.

In 15.3% of extrusions, extensive surface PS was found on extrusion membranes, appearing as PS rings surrounding the extrusion membrane (Fig 2Q). Since PS is typically found on the inner leaflet of the host plasma membrane, this subpopulation likely represents extrusions with permeabilized outer membranes. 24.4% of extrusions exhibited no surface PS (Fig 2Q). A similar punctate distribution of PS on the surface of extrusions was observed when extrusions were labeled *in situ*—immediately upon release into the supernatants of infected monolayers (data not shown). Little PS exposure was observed on the surface of HeLa cells infected with GFP expressing *C. trachomatis* for 24 h and 48 h (data not shown), consistent with data described previously [18].

Historical data suggests that mitochondria may localize in proximity to *Chlamydia* inclusions [19], and therefore were strong candidates to potentially associate with extrusions. We probed live extrusions with a mitochondria-specific dye and determined that 20.3% of extrusions contained mitochondria, with localizations at the extrusion periphery (Fig 2E-H, Q). The number of mitochondria in extrusions was small, ~1–5, even in those extrusions which contained them (Fig 2E-H). Fig 2I-L shows mitochondria staining in live, uninfected HeLa cells, confirming accurate staining of mitochondria. HeLa cells infected with mKate2-expressing L2 were also probed at 48 hpi (Fig 2M-P), and revealed very little mitochondria staining in infected cells. This may be due to the late stage of infection and the large size of the chlamydial inclusion, making mitochondria harder to spot. Mitochondria staining was also performed on early stage infected cells (18 hpi), which revealed abundant mitochondria staining in all areas of cytoplasm within the infected host cell (Suppl Fig 2). We additionally probed for the presence of the Golgi apparatus in extrusions; and like nuclei, this organelle was never observed (data not shown).

Fluorescence microscopy data of live extrusions therefore revealed that extrusions were largely devoid of host organelles, with the exception of occasional mitochondria. Transmission electron microscopy (TEM) of isolated extrusions confirmed the absence of mitochondria, Golgi apparatus and other discernible organelles within extrusions (Fig 2R). TEM images also showed that extrusions contained *C. trachomatis* encased by a remnant of the chlamydial inclusion, as shown by the presence of the extrusion’s double membrane (Fig 2R, inset). Consistent with other TEM images, the extrusion image in Figure 2R shows a distribution of chlamydial inclusion on the left portion of the extrusion (marked with an ‘I’ in Fig 2R), and host cell cytoplasm collected on the right half portion of the extrusion (marked with a ‘C’). This distribution is very similar to what was reported for *in vivo* sections containing *C. pecorum* extrusions [20]. It is possible that this unique separation of host cell cytoplasm and chlamydial inclusion within the extrusions may be artifact from centrifuging the extrusions into pellet form for fixation, or other electron microscopy handling procedures. We also imaged late-stage *C. trachomatis*-infected HeLa cells by TEM, using identical procedures, and did not detect this type of inclusion distribution in cells (data not shown). Thus, this property appears to be unique to chlamydial extrusions and not infected cells. Extrusions are thus defined as unique pathogenic structures that contain *Chlamydia*, host cytoplasm, chlamydial inclusion membrane, and plasma membrane that frequently contain externalized PS.

### Extrusions enhance the extracellular survival of *Chlamydia*

We next investigated whether extrusions were capable of delivering infectious *C. trachomatis* to epithelial cells, to inform if extrusions are viable vehicles for facilitating the cell-to-cell spread and dissemination of *Chlamydia.* Isolated extrusions were used to infect HeLa cells for 1–2 h, rinsed, and cells were cultured for another 12–24 h to allow infection. Evaluation of inclusion formation in cells revealed many infected cells that contained inclusions, and that were of a size equivalent to cells infected with free *C. trachomatis* EB (Suppl Fig 1). Thus, extrusions are infectious, and can function as vectors for chlamydial cell-to-cell spread, through breakdown of their membranes, thereby promoting exposure of infectious EB onto epithelial cells. We never witnessed prematurely large inclusions in cells that would be indicative of epithelial phagocytosis of extrusions; rather, the exposure of *C. trachomatis* to epithelial cells occurred through breakdown of extrusion membranes and release of their *Chlamydia* contents.

Outside the nurturing host cell environment, *Chlamydia* rapidly lose viability—within minutes to hours. We rationalized that extrusions may provide important supportive benefits to *Chlamydia*, conceptually as an ‘inclusion-like’ environment for these bacteria to inhabit as they traverse the extracellular milieu on their way to infect new epithelial cells. We therefore tested whether extrusions were capable of transiently preserving *C. trachomatis* survival and EB infectivity during culture outside of host cells. Extrusions were collected from infected HeLa cells at 72 hpi, and incubated at 37°C in cell-free growth media for up to 24 h. Equivalent suspensions of extrusions that were briefly sonicated to disrupt extrusion membranes, yielding an equivalent dose of *C. trachomatis* EB (‘free *Chlamydia*‘), were used as a control. At indicated time points, the infectivity of *C. trachomatis* from both groups (extrusions and free *Chlamydia*) was determined by gently sonicating samples and titering infectious bacteria onto McCoy cells for inclusion forming units (IFU) determination. Free *C. trachomatis* lost viability rapidly, with only 40% viable following 4 h incubation at 37°C, and 3% of original infectivity present after 24 h (Fig 3). Extrusion-derived *C. trachomatis* lost viability at a much slower rate; 76% viable EB were obtained from extrusions after 4 h at 37°C, and 32% infectivity of the extrusion-derived EB were obtained after 24 h (Fig 3). This disparity in survival of free vs. extrusion-derived EB at early time points may be the consequence of extrusions remaining largely intact up to 4 hpe (hours post extrusion). As time progressed, this gap lessened, which likely suggests that extrusions began to break down 4 h after their isolation. These results suggest that extrusion and its double membrane structure provides a niche environment, that may more closely mimic intracellular life within an inclusion, to allow enhanced survival of *Chlamydia* and protection from extracellular stresses.

**Fig 3.**
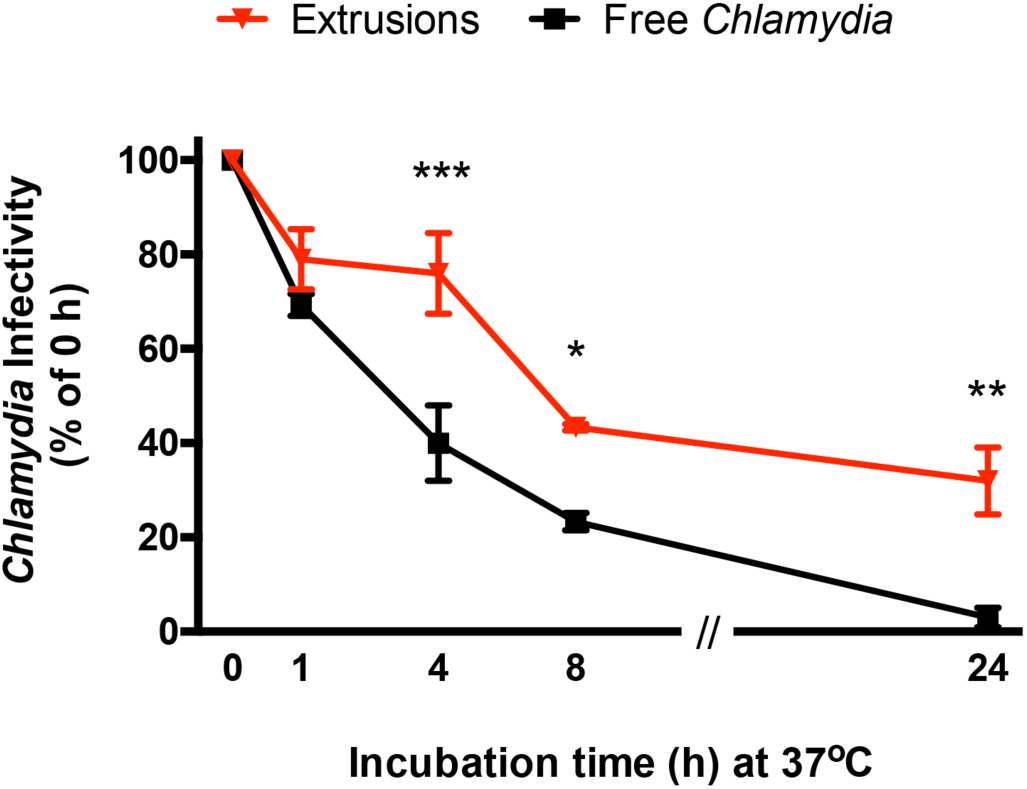
Extrusion-derived *Chlamydia* have prolonged infectivity outside of host cells. Intact extrusions or free *Chlamydia* were incubated in growth media at 37°C for times indicated, after which *Chlamydia* viability in samples was determined by brief sonication and direct infection onto McCoy cells. IFU were quantitatively determined for each sample, and data are expressed as a ratio of the t = 0 sample for each group. Data points show mean ± SEM, n ≥ 3. *** denotes a p value < 0.001, ** denotes a p value < 0.01, * denotes a p value < 0.05.

### Composition of developmental forms within extrusions remains unchanged

We postulated that one mechanistic basis for how extrusions could preserve *Chlamydia* infectivity is by facilitating continued RB–EB conversion, even while these bacteria are no longer host cell associated. The conversion of *Chlamydia* RB to infectious EB requires bacteria to be in an inclusion and host cell, may be due to contact of *Chlamydia* with the inclusion membrane [21,22], and therefore would not be expected to occur for *Chlamydia* released by lysis of the host cell.

To experimentally investigate whether the microenvironment of extrusions was sufficient to enable continued RB-EB conversion, we developed a high-magnification fluorescence microscopy strategy to accurately quantify distinct populations of *Chlamydia* RB and EB from extrusion samples. Extrusions were collected and incubated at 37°C from 0 to 8 h, and at specific time points samples were gently sonicated to break apart extrusions and evenly disperse the bacteria. Extrusion contents were subsequently plated onto coated coverglass, fixed, and labeled with a specific antibody to *C. trachomatis*. Random fields from all samples were imaged by fluorescence microscopy at 100× magnification to visually distinguish RB and EB within the population (Fig 4A). Computational imaging algorithms were designed to accurately identify *C. trachomatis* in images, and separate them into distinct populations of RB and EB (Fig 4A). The fidelity of measurement parameters was determined in advance using *C. trachomatis* preparations from sonicated 24 and 48 hpi infected cells (Fig 4B, left two columns), which were enriched for RB and EB, respectively. Data obtained from 24 hpi inclusions demonstrated an RB composition of 66%, whereas at 48 hpi inclusions were comprised of 15% RB and 85% EB (Fig 4B). There was a significant increase in EB percentage from 24 to 48 hpi, which was expected, due to the formation of EB toward the end of the *Chlamydia* developmental cycle.

**Fig 4.**
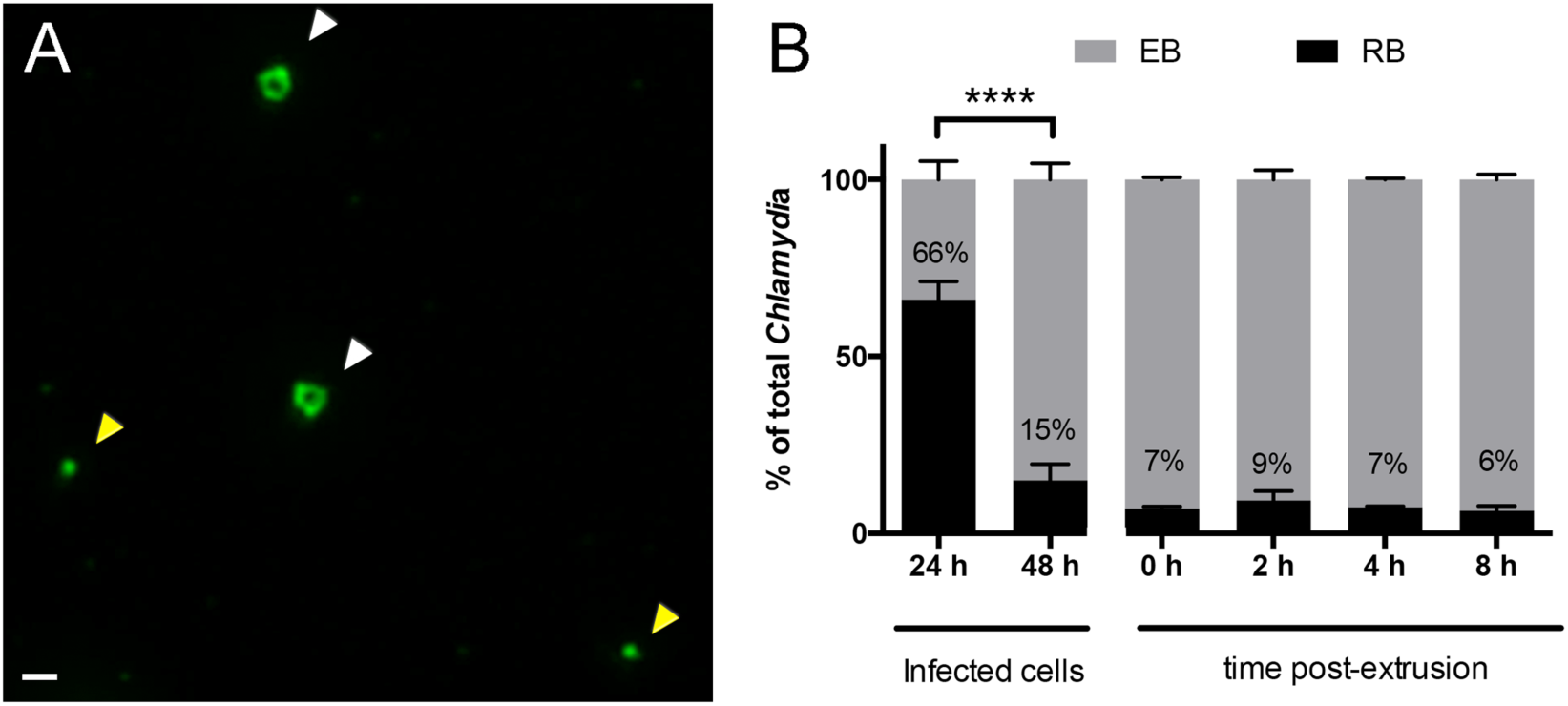
Ratio of chlamydial developmental forms within extrusions remain unchanged after release from host cells. *C. trachomatis* L2 extrusions were isolated and enriched from infected HeLa cells at 48 hpi, and incubated in cell culture media for 0 to 8 h at 37°C. At times indicated, extrusions were broken by brief sonication, and bacteria were plated onto poly-D lysine coated coverglass for fixation and immunofluorescence staining. *Chlamydia* were stained with an anti-MOMP antibody (green) and visualized by fluorescence microscopy at 100× magnification for illumination and differentiation of *C. trachomatis* RB and EB developmental forms. **A** Representative image of ringlike RB (marked with white arrow) and smaller EB (marked with yellow arrow). Scale bar, 1 μm. **B** Quantitative analysis of the relative ratio of EB and RB from 24 h and 48 h infected cells (left two columns), and experimental extrusion samples, (right four columns). Ratios were calculated from number of total bacteria per field per time group. Labels inside bars represent percentages of RB. Data show mean percentages ± SEM, n = 3. **** denotes a p value < 0.0001.

The developmental body composition of *C. trachomatis* taken from extrusions revealed a mean composition of 7% RB and 93% EB immediately after extrusions were collected from infected cultures, at 0 h. No significant increase in the percentage of EB was detected in extrusions from 0 h to 8 h at 37°C (Fig 4B, right 4 columns). Likewise, there was also no significant decrease in relative percentage of RB over the course of the 8 h incubation. These data indicate that *C. trachomatis* RB–EB conversion did not noticeably occur in extrusions, or else happened at a level below the threshold of detection for this assay, and suggests that RB–EB conversion requires more than just physical contact of *Chlamydia* with the inclusion membrane. Additional requirements for this key developmental transition likely include nutrients or energy, which extrusions either lack or only have in a finite supply compared to the parental host cell.

### Engulfment of *Chlamydia* extrusions by macrophages

The presence of externalized PS on the outer surface of extrusion membranes raised the possibility that *C. trachomatis* extrusions, unlike their malaria counterparts, might be readily recognized as apoptotic bodies by professional phagocytes, such as macrophages. We explored this possibility by infecting primary bone marrow-derived murine macrophages with chlamydial extrusions (derived from infections with GFP-expressing *C. trachomatis*), and the uptake of extrusions was determined by immunofluorescence analysis. Parallel samples of extrusion preparations were subjected to brief sonication to break up extrusion membranes, to result in freely dispersed *C. trachomatis* for infection of macrophages as a control. After early stages of incubation (0–4 h), intact extrusions were frequently found inside macrophages (Fig 5A), and rarely outside of macrophages. The size of *Chlamydia* clusters within macrophages at < 4 h far exceeded what would be expected from a nascent infection, and these clusters were never found in macrophages that were inoculated with sonicated extrusions (Fig 5D) or high MOI of *C. trachomatis* EB (data not shown). Thus, it can be concluded that these objects resulted from the engulfment of extrusions by macrophages. On average, extrusions were found inside approximately 5% of confluent macrophages for any given experiment, and incubated onto macrophages at a dose of ~0.1 extrusions for every macrophage. Similar fates were found for extrusions derived from other cell lines, of mouse or human origin, including L929, HeLa, and McCoy cell lines (data not shown). Extrusions were rarely imaged outside of macrophages or in mid-engulfment, and this is likely due to uptake occurring within the 1 h window of infection prior to immunofluorescence imaging.

**Fig 5.**
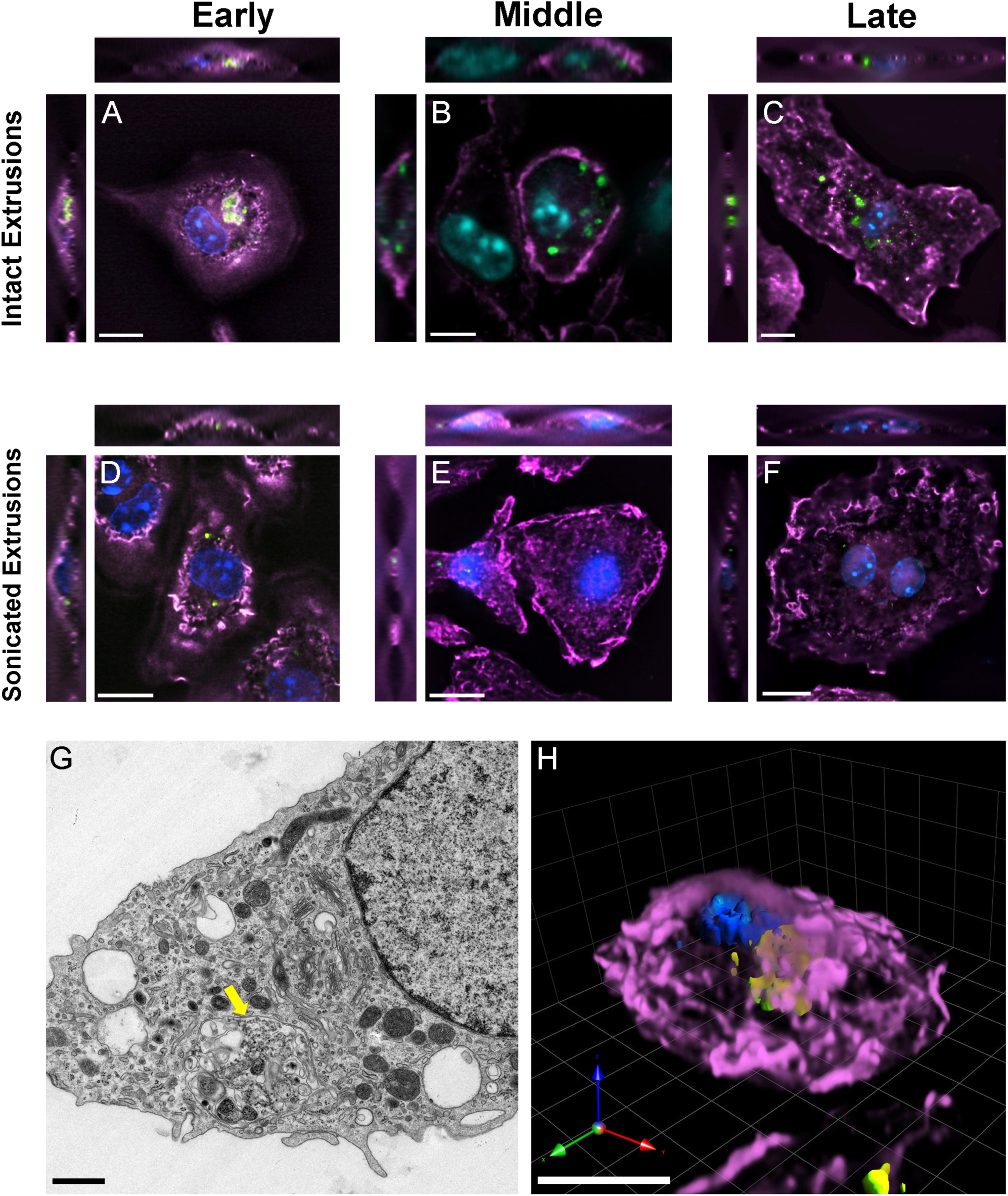
Engulfment of *Chlamydia* extrusions by primary macrophages. **Top row:** Isolated *Chlamydia* extrusions were incubated with bone-marrow derived macrophages for 1 h, then rinsed, and fresh macrophage media was plated onto cells for duration of assay; 0–72 h at 37°C. At stages indicated, cells were fixed and stained to visualize: GFP-expressing *C. trachomatis* (green), macrophage nuclei (DAPI, blue), and actin (phalloidin-647 purple). Representative images of macrophages containing *C. trachomatis* extrusions show distinct stages of their interaction: **A** early, 0–4 h, presence of intact extrusions; **B** middle, 4–24 hpi, broken-down extrusions, with *Chlamydia* still visible within macrophage; **C** late, 48–72 hpi, no visibly intact extrusions, but *Chlamydia* still visible within macrophage. In other experiments, isolated extrusions were briefly sonicated to release *Chlamydia*, and bacteria were incubated with macrophages for 0–72 h at 37°C. Representative examples are shown at similar stages as for extrusions: **D** early, some bacteria present in macrophages; **E** middle, very few to no bacteria seen within macrophage; **F** late, no bacteria seen in macrophages. **G** Transmission electron micrograph of a representative macrophage containing an engulfed *C. trachomatis* extrusion following 1 h co-incubation (shown by yellow arrow). Scale bar, 1 μm. **H** Three dimensional view of a macrophage containing an engulfed extrusion. Host cell actin (purple), nuclei (blue), and GFP *Chlamydia* (GFP). Scale bar, 10 μm for all panels except **G**.

At later times post-engulfment, 48–72 h, GFP-*C. trachomatis* from broken-down extrusions were commonly seen dispersed within large areas inside the macrophages (Fig 5B-C). Although engulfed *C. trachomatis* from sonicated extrusions were readily visible at early stages of infection, they were altogether cleared from macrophages at these later times (Fig 5E-F). For macrophages that had engulfed extrusions, no gross morphological changes were apparent; in some instances, macrophages harboring extrusions appeared more ‘rounded’ than uninfected cells, or macrophages containing phagocytosed free *C. trachomatis*.

TEM was performed to confirm the presence of intact extrusions within macrophages at 1 h post incubation. In Figure 5G, an intact extrusion can be seen within the cytoplasm of the macrophage, shown by the yellow arrow. Three-dimensional rendering of a fluorescently-labeled macrophage, containing an intact extrusion following a 1 h co-incubation, was also performed (Fig 5H). In this example, host cell actin completely covered the large, green extrusion, confirming engulfment by the macrophage.

Other rare phenotypes were occasionally observed in macrophages after engulfment of extrusions. In some instances, advanced stages of culturing revealed some instances of macrophages containing fully-formed *C. trachomatis* inclusions (Suppl Fig 3A). We confirmed that these vacuoles were inclusions, and not long-lasting extrusions, by staining cells with an antibody to the chlamydial inclusion protein CT223 (Suppl Fig 3A). A punctate ring staining pattern for CT223 was unique to these rare examples, as the presence of CT223 was never detected in macrophages containing intact, engulfed extrusions at early times of infection, due to the subsequent breakdown of extrusion membranes by the macrophage (Suppl Fig 3B). Additionally, extrusions with enlarged *Chlamydia*, resembling aberrant bodies, were found (Suppl Fig 3C). This suggests that RB contained in extrusions may be capable of arrested development. In rare cases in the wells of macrophages infected with free *C. trachomatis*, bacteria could be seen in late stage infection (Fig S3D), and it is possible that a small population of free *C. trachomatis* were able to survive and form inclusions within macrophages, even though no inclusions were ever seen from this infection group. This result is consistent with previous literature that demonstrated an inability of *C. trachomatis* to capably infect macrophages [23].

### Extrusion-derived *Chlamydia* avoid killing by macrophages

Because *C. trachomatis* were routinely present inside macrophages that had engulfed extrusions compared to free bacteria, and at late times post-engulfment (e.g., >24 h), we hypothesized that extrusion-derived *C. trachomatis* may benefit from preferential survival inside phagocytic cells. To test this hypothesis, extrusions and free *C. trachomatis* of the same dose were infected onto macrophages for 1 h, rinsed, then incubated for 8 h at 37°C. For each macrophage experiment, approximately 2.5 × 104 extrusions were incubated onto 3 × 105 macrophages, corresponding to roughly 0.1 extrusions per macrophage. It is important to note, however, that the same dose of extrusions onto McCoy cells for IFU analysis corresponded to an MOI ~1. Following the 1 h incubation, macrophages were lysed to release bacteria, and *Chlamydia* infectivity was quantitatively measured by titering EB onto McCoy cells for IFU determination. From macrophages infected with free *C. trachomatis*, EB were only recoverable at 1 h (6.2 IFU/field at 20× magnification), with no IFU detected at 4 or 8 h (Fig 6). In contrast, significantly more infectious EB were recoverable from extrusion-containing macrophages at 1 h (25.4 IFU/field), and even out to 8 h. These data indicate that extrusions were able to transiently protect *C. trachomatis* from phagolysosomal killing by macrophages. The functional consequence of extrusion-derived chlamydial survival inside macrophages for up to 8 h, and to a lesser extent even 24 h (data not shown), is unclear. One expected *in vivo* outcome would be migration of these macrophages across tissues or to lymph nodes, potentially to elicit differential immune pathway activation and cytokine secretion.

**Fig 6.**
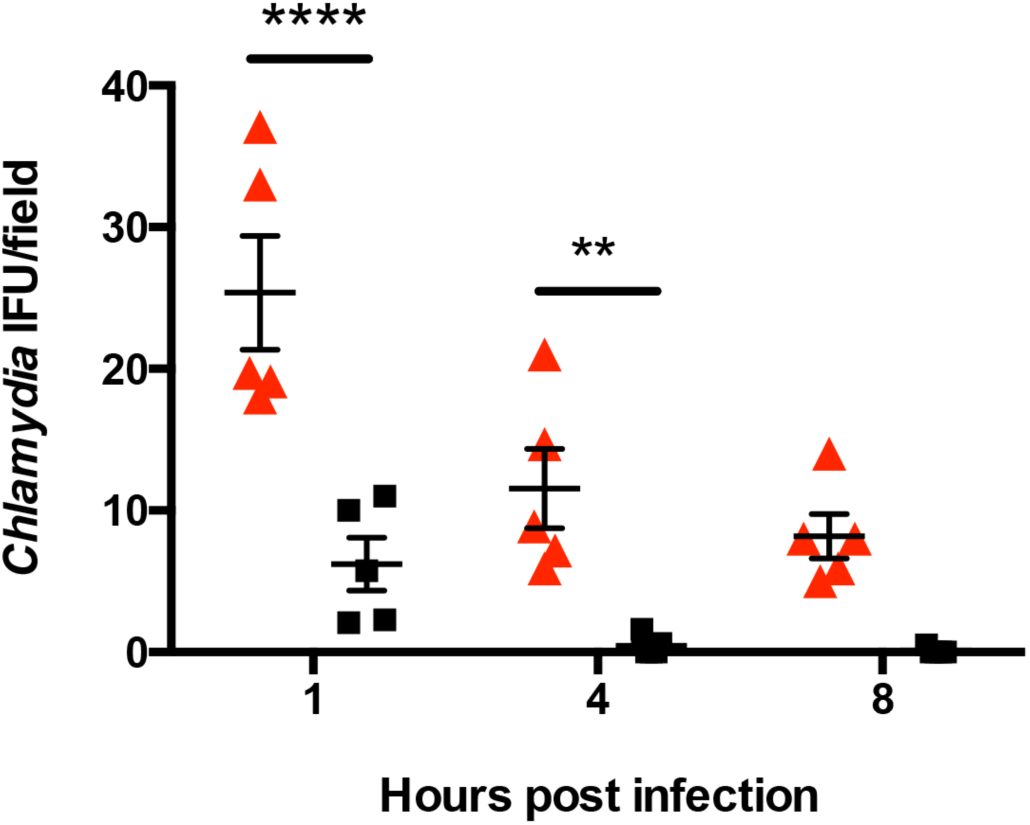
Preferential survival of extrusion-encased *Chlamydia* within primary macrophages. Extrusions, or sonicated extrusions (free *Chlamydia*), were incubated onto murine bone marrow derived macrophages and incubated at 37°C for up to 8 h. At times indicated, cells were sonicated to release bacteria, and *C. trachomatis* infectivity was quantitatively measured by performing IFU assays on McCoy cells. Data points show mean ± SEM, n = 3. **** denotes a p value < 0.0001, ** denotes a p value < 0.01.

### Escape and infectivity of extrusion-derived *Chlamydia* from macrophages

The discovery that extrusion-derived *C. trachomatis* were capable of surviving inside macrophages raised intriguing questions about the beneficial outcomes *Chlamydia* might gain from this phenomenon. One possibility is that chlamydial extrusions induce eventual death of the host macrophage, thereby releasing *Chlamydia* into the extracellular milieu. If macrophage killing and *Chlamydia* escape were to occur after macrophage migration, it could provide an opportunity for *C. trachomatis* to disseminate in the female genital tract (i.e., ascend to upper genital tract tissue), or evade clearance by innate immune cells. To test whether extrusion-derived *C. trachomatis* were capable of eventually escaping the macrophage, we infected primary macrophages with extrusions, or sonicated extrusions, (free *Chlamydia*) for 2 h (Fig 7A). Cells were rinsed fully and incubated at 37°C for up to 48 h. Daily rinses of cultures were performed to ensure no carryover of infectious *C. trachomatis* that had escaped prior to the indicated time points. At 0 and 48 hpi, supernatants were collected from macrophages and the quantitative burden of infectious *C. trachomatis* EB in supernatants was determined by IFU titering onto McCoy cells. Immediately after infection and rinsing, less than 1 IFU per field at 20X magnification of *C. trachomatis* EB were recoverable from macrophages (Fig 7B), indicating that most extracellular bacteria and extrusions had either been engulfed by macrophages or efficiently removed by rinsing. A significant amount of extrusion-derived EB (59.3 IFU/field at 20× magnification) were present in macrophage supernatants at 48 h after extrusion engulfment (Fig 7B). This was in marked contrast to macrophages that were infected directly with free *C. trachomatis* EB; for these cells only a minimal amount of viable bacteria (3 IFU/field) were present in culture supernatants at 48 h (Fig 7B). We postulate that chlamydia are able to exit the macrophage through host cell death, which has been observed after late times post-infection with extrusions (data not shown). Infection of macrophages with free *C. trachomatis* did not show the same cell death in macrophages. These results provide compelling evidence that unlike phagocytosed free *C. trachomatis*, extrusions provide a temporary niche for the transient survival and passage of *Chlamydia* inside primary macrophages. Furthermore, this trafficking strategy does not lead to a ‘dead end’ for *C. trachomatis*; after avoidance of phagocytic killing, these extrusion-derived *Chlamydia* are able to ultimately leave the macrophage and infect new epithelial cells.

**Fig 7.**
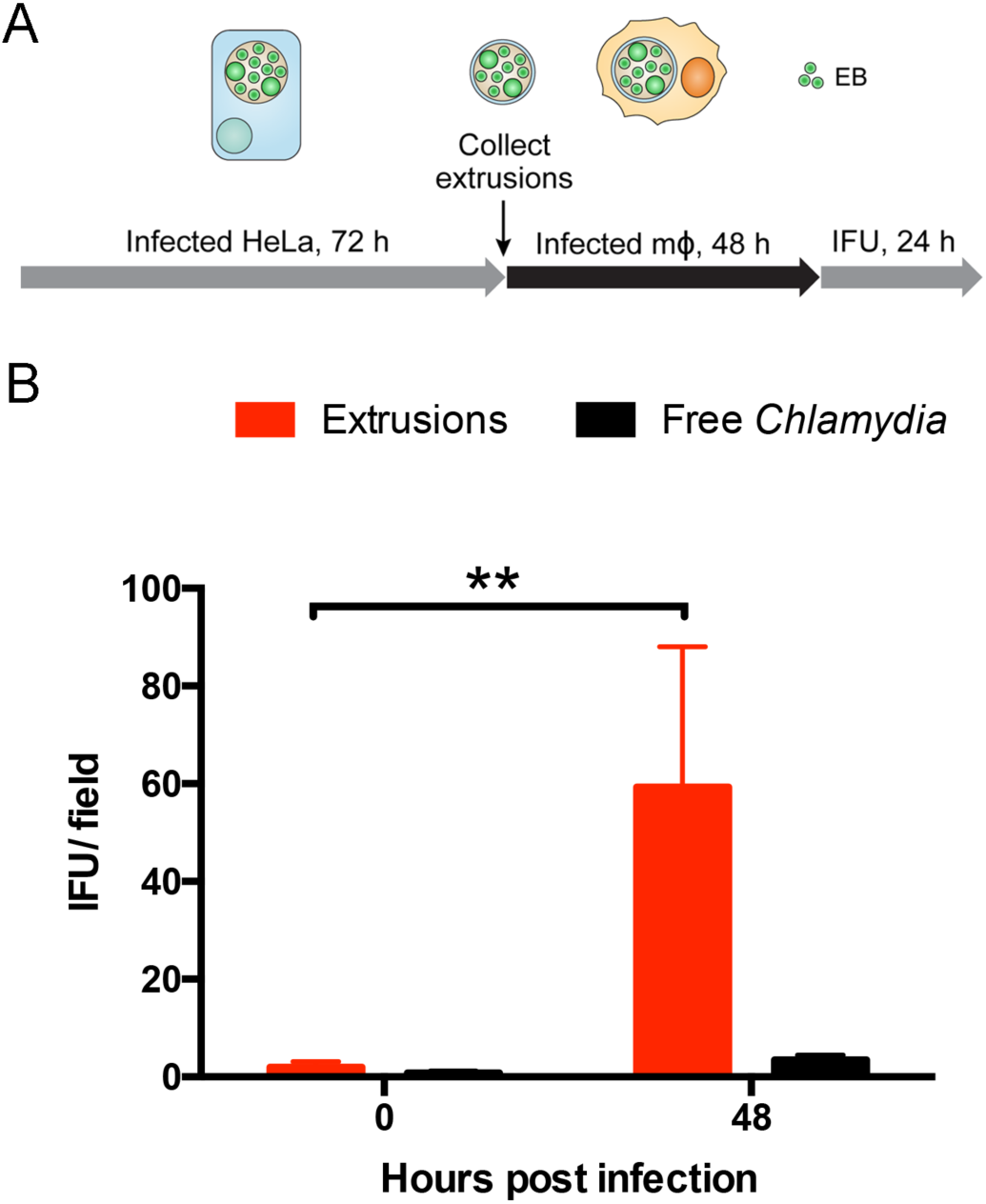
Release of *Chlamydia* EB from extrusion-infected macrophages. A Overview of the experimental strategy depicting extrusion isolation, infection times in macrophages, and collection of released *C. trachomatis* EB. **B** Extrusions, or sonicated extrusions, were used to infect macrophages, and cell supernatants were collected immediately after rinsing (t = 0) and at 48 h. The infectivity of *Chlamydia* in macrophage supernatants was determined by IFU assays on HeLa cells, at 24 h. Data points show mean ± SEM, n = 3. ** denotes a p value < 0.01.

## Discussion

Since the discovery of extrusion as a *Chlamydia* exit mechanism, its functional role and contribution to chlamydial pathogenesis remains unknown. The engulfment and transient persistence of the chlamydial extrusion within a macrophage represents a novel discovery for bacterial pathogenesis (Fig 8). Early after internalization, extrusions existed as intact structures containing dozens to hundreds of bacteria. Within hours, most extrusions began to dissolve and bacteria were routinely found dispersed throughout the macrophage, and not in a vacuole. Coincident with this progression, a significant percentage of extrusion-derived *C. trachomatis* were able to withstand intracellular killing by macrophages, as evidenced by the continued (yet declining) presence of infectious *C. trachomatis* EB at all times of their residence in macrophages. This was in stark contrast to the fate of phagocytosed free *C. trachomatis* by macrophages. Consistent with the literature [23], engulfed free bacteria were rapidly killed by primary macrophages, and only a small percentage were capable of establishing an infection.

**Fig 8.**
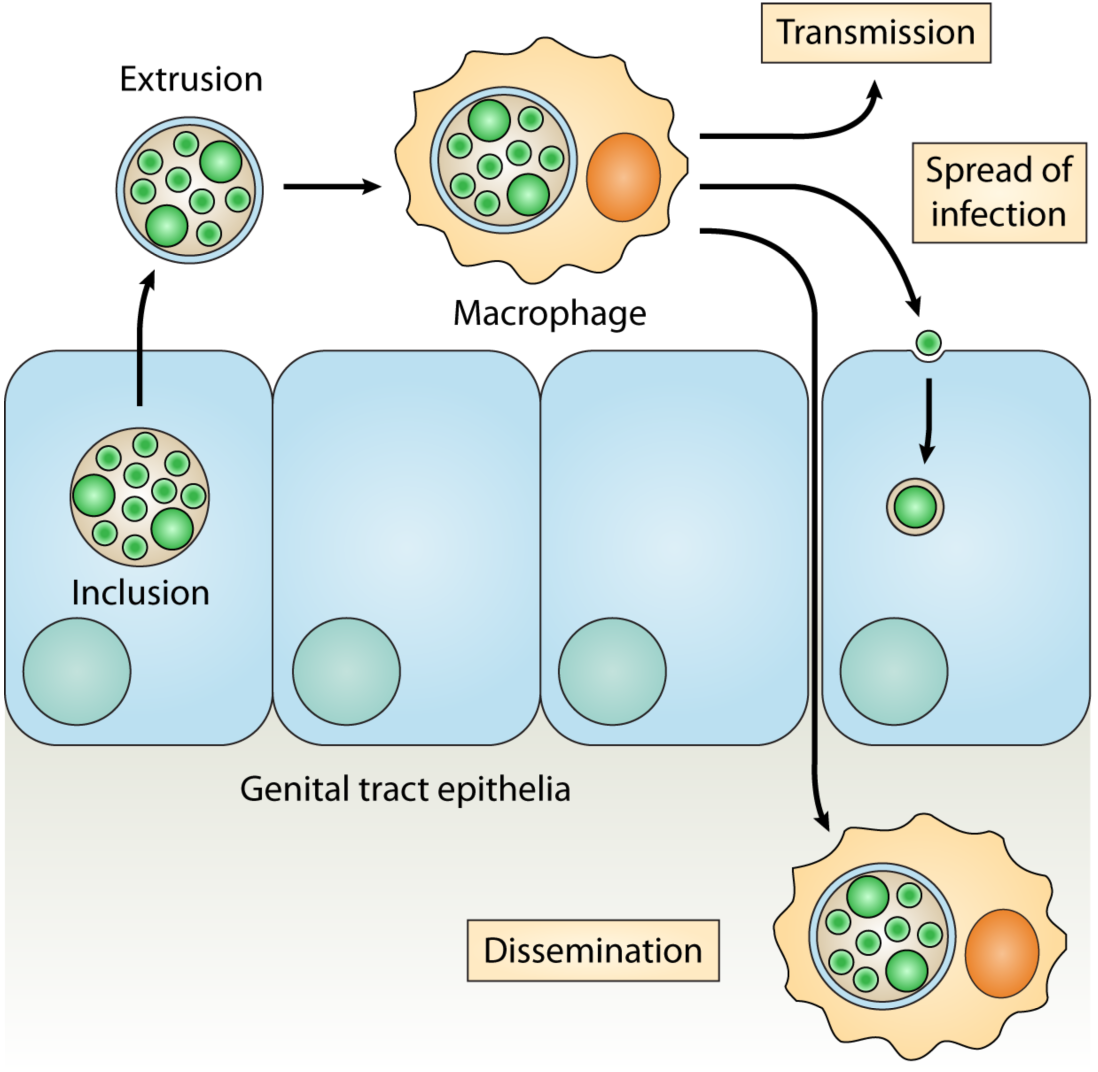
Model for the role of extrusions and macrophages in the dissemination of *C. trachomatis* in the upper female genital tract. Upon release from infected epithelial cells, *Chlamydia*-containing extrusions are engulfed by macrophages. Migration of these macrophages, followed by eventual escape of *Chlamydia* from them, can result in the dissemination of infectious *C. trachomatis* to more distant sites, e.g., away from inflammatory foci surrounding the primary site of infection, to draining lymph nodes, or to new hosts. Extrusions may alternatively mediate some of these outcomes without the need for hijacking macrophages.

At late stages of extrusion-derived *C. trachomatis* survival in macrophages, infectious bacteria ultimately emerged in macrophage supernatants, suggesting that these *Chlamydia*, or undefined factor(s) contained within extrusions, can lead to macrophage lysis. We do not think extrusions can form from macrophages that have engulfed extrusions, since no examples were ever detected. We hypothesize that after their breakdown inside macrophages, extrusions present a high number of *C. trachomatis* and *Chlamydia*-derived pathogen-associated molecular pattern(s) (PAMPs) to cytoplasmic PAMP sensing pathways in the macrophage that ultimately trigger macrophage lysis. Extrusions deliver a higher burden of *C. trachomatis* to the cytoplasm of macrophages than we have can achieve artificially by internalization of high MOI of bacteria. We are currently working on defining the unique cellular responses of macrophages to chlamydial extrusions, as it carries important consequences for the host response to *C. trachomatis* infection *in vivo*.

In the course of empirically deriving a simple experimental approach for isolating and enriching endogenously produced extrusions from *Chlamydia* infected cells, we exhaustively attempted many alternative approaches, but ultimately, the most efficient and reliable approach was to let the infected monolayers produce and release chlamydial extrusions into culture supernatants. Performing this collection in fresh growth media or saline, for a defined window of time, resulted in the best balance of extrusion quality and yield. We determined that extrusion quality was an important factor and goal for downstream functional studies using extrusions. Although chlamydial extrusions were durable enough to withstand successive rounds of centrifugation, they intrinsically permeabilized over time. No deviations in the collection and handling strategy were able to completely prevent this phenomenon, for example supplemental bovine serum albumin [24], or the addition of protease inhibitors [8]; however, our derived approach resulted in the least amount of breakdown among procedures tested. Collectively, these findings point to an innate tendency for extrusion membranes to weaken and permeabilize. As it pertains to the natural history of *C. trachomatis* infections, this outcome is probably a beneficial one for the bacteria. In this manner, extrusions can provide a temporary, protective niche for extracellular *Chlamydia* while also ensuring that *Chlamydia* ultimately escape to spread to new epithelial cells.

The successful development of a method for isolating *Chlamydia* extrusions from host cells enabled the detailed examination of the internal composition of these structures. Our data demonstrate that the chlamydial extrusion is essentially an extracellular, pathogenic compartment. With the exception of occasional mitochondria, extrusions were largely devoid of host cell organelles. This indicates that the mechanism of extrusion formation is not due to wholesale division or cleavage of the infected host cell—wherein half of the cell gets released as an extrusion along with whatever host cell organelles that happened to accompany it. Instead, the extrusion mechanism appears to have evolved to produce a compartment of *Chlamydia* that are encased by vacuole and host plasma membranes, and little more. One can speculate that the small numbers of mitochondria that occasionally, albeit rarely, accompany extrusions may provide a temporary source of energy or oxidative buffering for these extracellular *Chlamydia*.

Interestingly, phosphatidylserine externalization was commonly associated with extrusion outer membranes. An important functional outcome of PS externalization on extrusion surfaces is that these objects mimic a critical component of apoptotic bodies for promoting their uptake by phagocytic cells, such as macrophages and dendritic cells. *Chlamydia* extrusions were efficiently engulfed by macrophages, to an extent where they were easy to identify in macrophage monolayers. In experiments not shown, we have discovered that cells infected with urogenital, LGV, and ocular strains of *C. trachomatis*, *C. pneumoniae*, *C. muridarum*, and *C. psittaci*, all produce extrusions. Furthermore, all of these extrusions are engulfed by macrophages and dendritic cells, and with extrusions derived from either human (HeLa) or murine (L929, McCoy) cells.

Although *Chlamydia* extrusions represent unique structures in the bacterial world, they share morphological similarities with merosomes that are produced by *Plasmodium* spp. [14-17]. After their liver stage conversion from sporozoites to merozoites, *Plasmodium* exit hepatocytes in membrane-bound compartments containing thousands of parasites. Unlike *Chlamydia* extrusions, *Plasmodium* merosomes suppress externalization of PS, and it has been shown that this allows malarial parasites to evade interactions with macrophages and efficiently transition to the blood stage of malaria infection [14-16]. It is fascinating to consider both the convergent evolution of the extrusion-like structure for exiting host cells, and also its variable roles for disseminating within a host.

The results of this work raise intriguing hypotheses and questions regarding the evolutionary development of the extrusion strategy by chlamydiae. Our data demonstrate tangible benefits for *Chlamydia* that exit epithelial cells within extrusions. These bacteria are able to survive in the extracellular environment much longer than free *C. trachomatis* EB. The membranous confines of extrusions provide intracellular-like benefits to *Chlamydia* even while they navigate the extracellular environment. These benefits may include enhancing retention of nutrients, energy, or other factors vital to chlamydial viability maintenance of EB infectivity. In this manner, we view the *Chlamydia* extrusion as a transient, extracellular niche, from which *Chlamydia* can break out to infect new epithelial cells—likely through delivery of a bolus of infectious EB—and also be protected from extracellular immune defenses. It is conceivable that the release of ‘packages’ of EB onto nascent epithelial cells results in more efficient infection *in vivo*. Given the natural history of chlamydiae as obligate intracellular bacteria, their evolution of promoting an extracellular compartment that partially resembles their intracellular home makes sense teleologically. Moreover, our data suggest that a major outcome of chlamydial exit within extrusions is to facilitate their uptake by and survival within macrophages. We propose a model in which *C. trachomatis* exploits macrophages for the advantage of dissemination within a host (away from inflammatory foci), and potentially host–host transmission, as has been proposed for *Neisseria gonorrhoeae* [25] and *Staphylococcus aureus* [26].

## Methods

### Cell culture, *Chlamydia* propagation and infection

HeLa 229 and McCoy cells, were routinely grown in RPMI 1640 media supplemented with 10% FBS (HyClone, Thermo Fisher Scientific, Rockford, IL) and 2 mM L-glutamine (HyClone), at 37°C with CO2. For all microscopy experiments, cells were subcultured and plated onto chambered coverglass slides (Lab-Tek II; Nunc, Rochester, NY), or glass bottom culture dishes (MatTek, Ashland, MA) or 6-well and 24-well plates (BD Falcon).

*C. trachomatis* serovar L2 (LGV 434) was propagated in L929 cells grown in suspension culture or HeLa cells grown in T75 flasks and purified as previously described [27]. Chlamydial EB were isolated by sonic disruption of L929 suspensions and purification by centrifugation. The final L2 pellet was resuspended in sucrose phosphate buffer (SPG; 5 mM glutamine, 0.2 M sucrose. 0.2 M phosphate buffer) and stored at −80°C.

Primary bone-marrow derived murine macrophages were prepared and frozen from femurs of 6 to 8-week old C57BL/6 female mice (Jackson Laboratory, Bar Harbor, ME). Macrophages were grown in RPMI 1640 media supplemented with 20% FBS (HyClone, Thermo Fisher Scientific, Rockford, IL), 10% M-CSF, and 2 mM L-glutamine (HyClone), at 37°C with CO2. For all experiments, cells were subcultured and plated onto chambered coverglass slides (Lab-Tek II; Nunc, Rochester, NY), glass bottom culture dishes (MatTek, Ashland, MA) or 24-well plates (BD Falcon).

Infections were performed by washing cells with Hank’s balanced saline solution (HBSS, HyClone) and incubating cells with *Chlamydia* EB diluted in HBSS to a multiplicity of infection ≤1, for 2 h at 25°C. Following static incubation, cells were rinsed twice with HBSS, re-immersed in growth media and incubated at 37°C.

### Extrusion isolation

*C. trachomatis* serovar L2 was grown in semiconfluent HeLa cells for 72 h, or in McCoy cells for 48 h in RPMI supplemented with 10% FBS, L-glutamine and cycloheximide (2 μg/mL). Media on cell monolayers was removed, rinsed, and new media added to infected cultures at 72 hpi or 48 hpi (for McCoy cells). Infected cell cultures were allowed to proceed with infection in new media for 2–4 h to endogenously collect extrusions, then media was centrifuged at 75 × g for 5 m, followed by removal of supernatant and a second centrifugation spin at 1200 rpm for 5 m. The extrusion pellet was immediately resuspended in fresh RPMI. To enumerate the number of extrusions obtained from a cell monolayer, resuspended extrusions were stained with SYTOX Green (1:2000, Molecular Probes) and Hoechst (1:2000, Molecular Probes) for 5 minutes at 25°C, plated as 10–20 μl drops onto glass slides and imaged immediately on an inverted fluorescence microscope. Intact extrusions were identified as having chlamydial inclusions, lacking nuclei and being the appropriate size.

### Sonication of extrusions

Extrusions were collected using the method described above, and separated into two populations. One population was sonicated using a hand sonicator set at 20 A, and placed into a conical tube of extrusions for 2–3 seconds, 10 times, while on ice. This setting and brief sonication method was used to rupture the extrusion membranes to release free *Chlamydia*, and also keep the viability of the sonicated and extrusion population equal.

### Immunofluorescence and live fluorescence microscopy

All live microscopy and immunofluorescence was performed on a Nikon Eclipse Ti inverted fluorescence microscope. Image capturing performed using Hammamatsu camera controller C10600 and Volocity imaging software, version 6.3 (PerkinElmer; Waltham, MA). Infected cells were fixed in 3.7% paraformaldehyde (Ted Pella) 15 m, then permeabilized with 0.1% Triton-X-100 (Fisher), blocked with 1% BSA-PBS (Fisher), and stained. Antibodies/dyes were obtained from the following sources: Phalloidin 633, donkey anti-goat 488 from Invitrogen (Waltham, MA), DAPI, goat anti-mouse 488 from Thermo Fisher (Waltham, MA), anti-GFP 488, FM4–64 from Molecular Probes (Eugene, OR), MitoTracker Green, Annexin-V 568 from Life Technologies (Carlsbad, CA), mouse anti-*Chlamydia* FITC conjugate from Meridian Diagnostics (Cincinnati, OH), goat anti-*C. trachomatis* MOMP from Virostat (Portland, ME), mouse anti-*C. trachomatis* LPS ad mouse anti-CT223 donated by Bob Suchland (University of Washington, WA).

### Image processing and analysis

Three-dimensional image stacks were further processed in Volocity by performing illumination correction (in *z* dimension) and deconvolution (25 iterations). Individual *xy* and *xz* slices were obtained from image stacks for figure assembly. Three dimensional opacity renderings of fluorescent image stacks were generated in Volocity. Minor retouching of all micrographs—for example, color assignment, contrast adjustment, RGB merges and cropping—were performed with Volocity and Photoshop CS6 (Adobe). Photoshop CS6 (Adobe) was used to assemble all figures into their final form.

### Electron microscopy

Extrusions, or macrophages containing extrusions were collected and cold fixative added, and centrifuged to form tight pellet. Supernatant was removed and cold fixative was added on top of cell or extrusion pellet, and centrifuged once more. All samples were kept at 4°C. All samples were fixed in 2% glutaraldehyde, 1% paraformaldehyde in 0.1M sodium cacodylate buffer pH 7.4, post fixed in 2% osmium tetroxide in the same buffer, then block stained with 2% aqueous uranyl acetate, dehydrated in acetone, infiltrated, and embedded in LX-112 resin (Ladd Research Industries, Burlington, VT). Samples were ultrathin sectioned on a Reichert Ultracut S ultramicrotome and counter stained with 0.8% lead citrate. Grids were examined on a JEOL JEM-1230 transmission electron microscope (JEOL USA, Inc., Peabody, MA) and photographed with the Gatan Ultrascan 1000 digital camera (Gatan Inc., Warrendale, PA).

### Statistical analysis

Statistical evaluation of data was performed by calculating the standard error of the mean (SEM) or using a two-way analysis of variance (ANOVA). P values <0.05 (*) were considered statistically significant. P-values of <0.001 (**), <0.0001 (***), and <0.00001 (****) were marked as indicated. Calculations were performed in Prism (Graphpad) and Microsoft Excel.

## Acknowledgments

We thank Ashley Sherrid for assistance with isolating bone marrow derived macrophages, and for discussions regarding this study. We thank Richard Stephens for antibodies and intellectual discussions. We thank Jinny Wong at the UCSF Gladstone Electron Microscopy Core for assistance and guidance with electron microscopy. We thank Scott Hefty for his gift of mKate2-expressing *C. trachomatis* L2. We also thank Bob Suchland for antibodies and for discussions regarding this study.

## Author Contributions

Conceived and designed the experiments: MZ, KH. Performed the experiments: MZ, TE, AV Analyzed the data: MZ, KH. Generated figures and wrote the paper: MZ, KH. This work was supported by R01AI095603.

## Supporting Information

**Fig S1.**
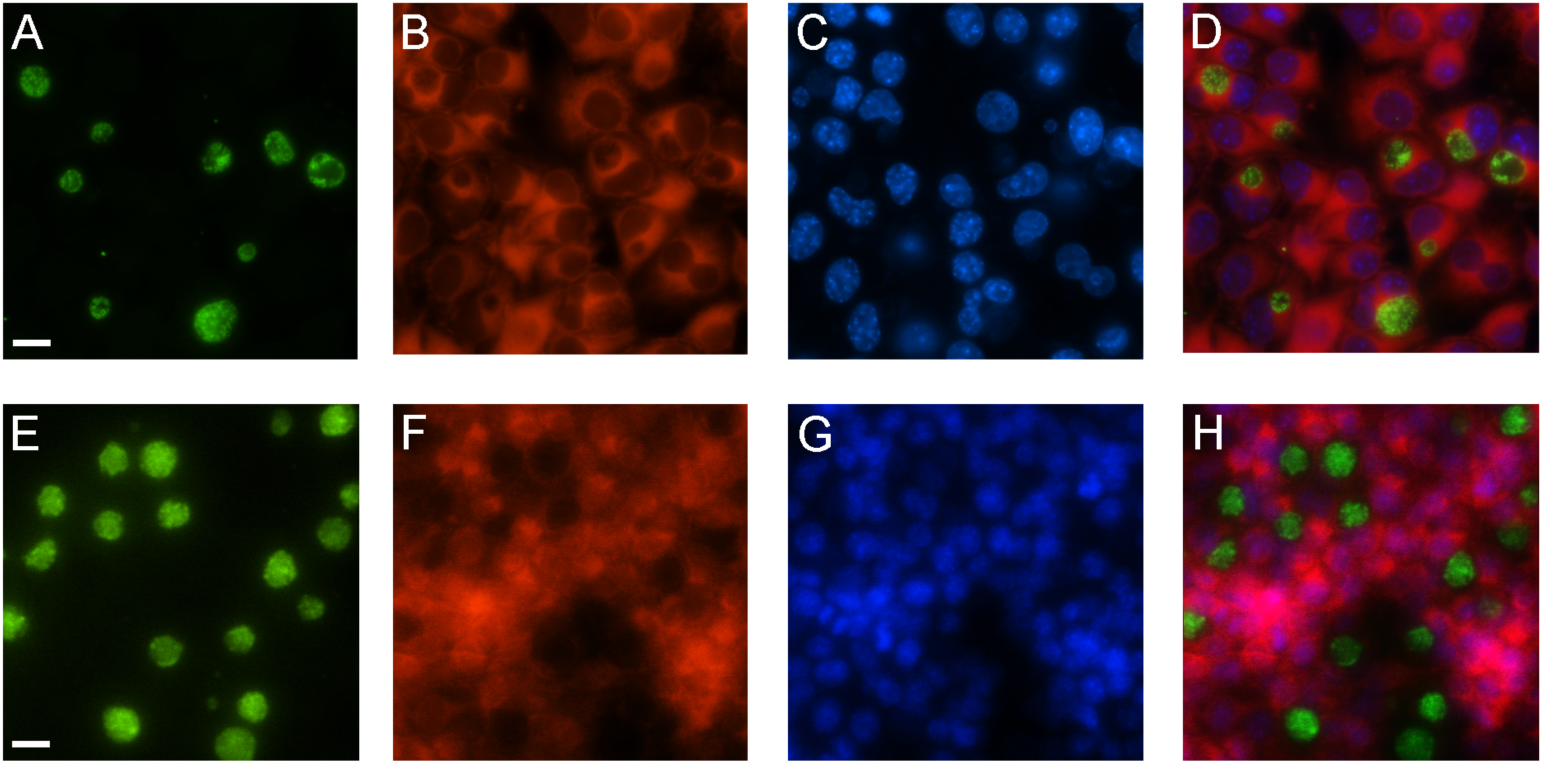
Extrusions are infectious, and not phagocytosed by HeLa cells. **Top row:** *C. trachomatis* extrusions were incubated onto HeLa cells for 2 h, then rinsed, and allowed to incubate at 37C for 18 hpi. Cells were fixed and stained for immunofluorescence analysis. **A** GFP-expressing *Chlamydia* (green). **B** Evans blue counterstain of HeLa cytoplasm (red). **C** DAPI staining of host cell nuclei (blue). **D** Merge of panels **A-C. Bottom row:** HeLa cells infected with GFP-*Chlamydia* 18 hpi. **E** GFP-expressing *Chlamydia* (green). **F** Evans blue counterstain of HeLa cytoplasm (red). **G** DAPI staining of host cell nuclei (blue). **H** Merge of panels **E-G**. Scale bar, 10 μm.

**Fig S2.**
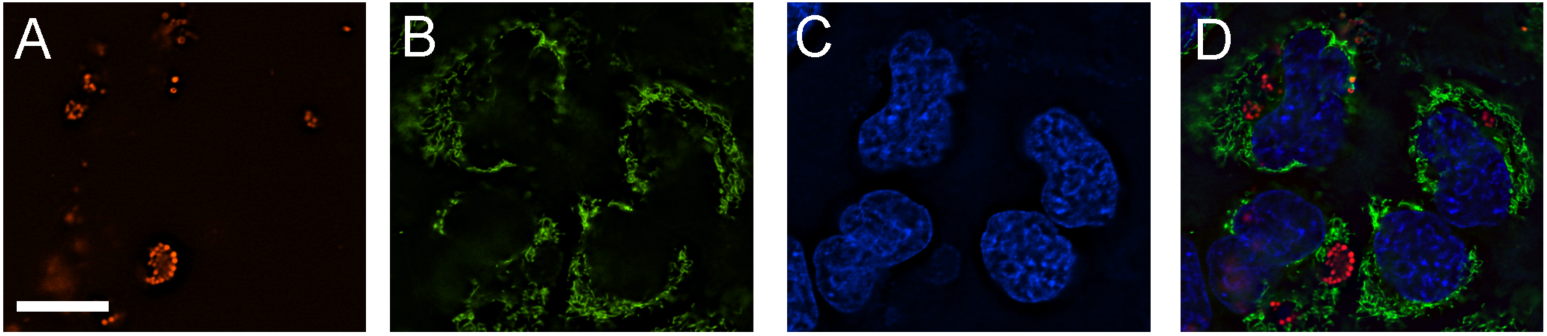
Immunofluorescence of early stage infected HeLa cells. HeLa cells were infected with mkate expressing *C. trachomatis* for 2 h, then rinsed and incubated at 37C for 18 hpi. **A** mKate2-expressing *C. trachomatis* (red). B Mitochondria stained with MitoTracker Green FM (green). **C** DAPI staining of host cell nuclei (blue). **D** Merge of panels **A-C**. Scale bar, 10 μm.

**Fig S3.**
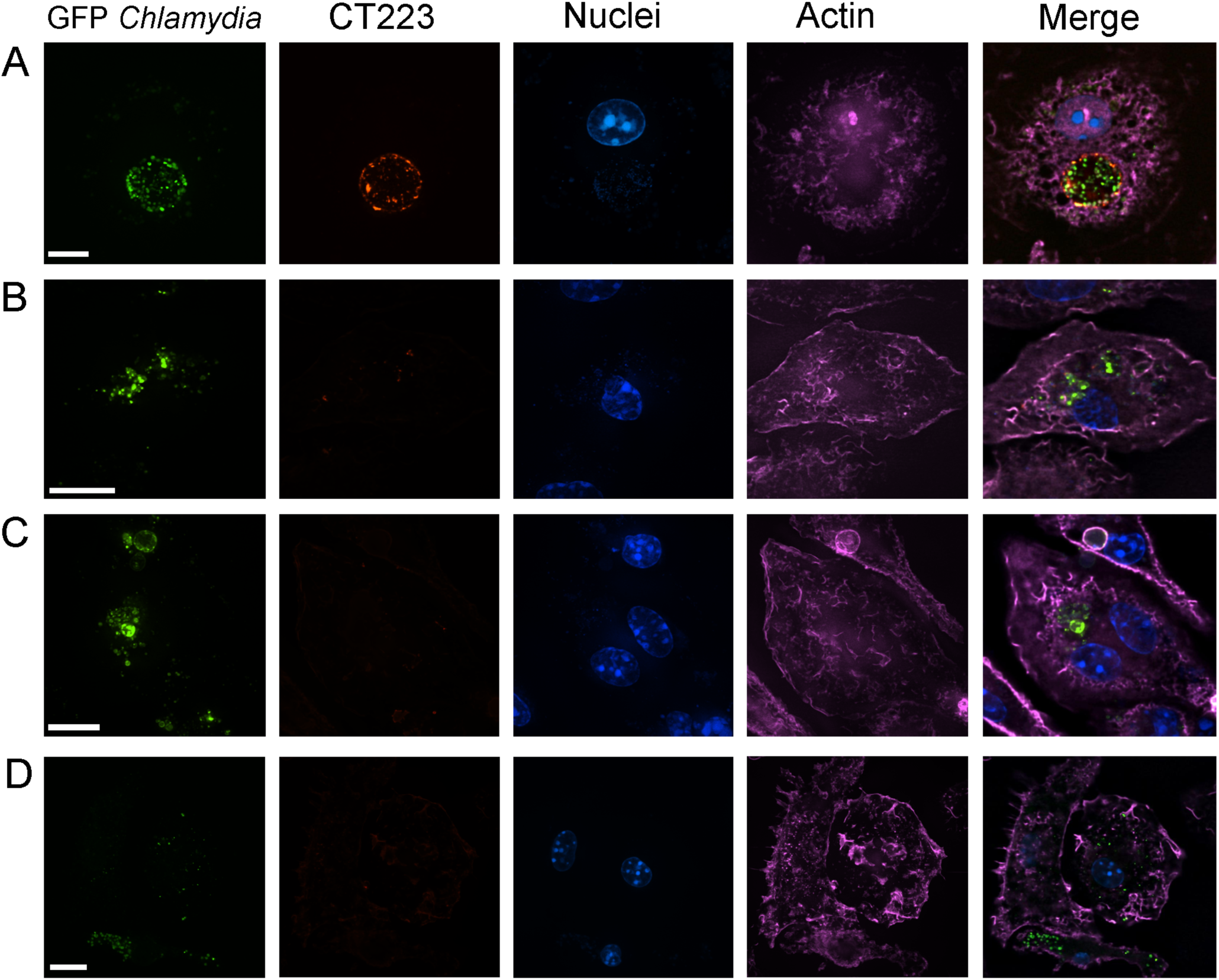
Additional immunofluorescence of murine macrophages, showing less prevalent phenotypes. **A** Murine macrophage incubated with extrusion at 24 hpi, showing establishment of an inclusion. **B** Macrophage incubated with extrusions at 0 hpi, showing no CT223 staining around the periphery of the inclusion. **C** Extrusion engulfment 0 hpi showing what appears to be a small abberant body within the macrophage. **D** Free *Chlamydia* engulfment by macrophage showing survival of small population of *Chlamydia* 24 hpi. For all images in figure: From left: GFP-expressing *C. trachomatis* (Green), inclusion membrane staining with antibody to CT223 (red), nuclei stained with DAPI (blue), actin staining with phalloidin-633 (purple), and merge of 4 colors. Scale bar, 10 μm.

## References

1. Burton MJ, Mabey DCW. The global burden of trachoma: a review. PLoS Negl Trop Dis. 2009;3: e460.

2. Gerbase AC, Rowley JT, Heymann DH, Berkley SF, Piot P. Global prevalence and incidence estimates of selected curable STDs. Sexually Transmitted Infections. 1998;74 Suppl 1: S12–6.

3. Darville T, Hiltke TJ. Pathogenesis of genital tract disease due to *Chlamydia trachomatis*. J Infect Dis. 2010;201 Suppl 2: S114–25.

4. Brunham RC, Rey-Ladino J. Immunology of Chlamydia infection: implications for a *Chlamydia trachomatis* vaccine. Nat Rev Immunol. 2005;5: 149–161.

5. Hafner LM, Wilson DP, Timms P. Development status and future prospects for a vaccine against *Chlamydia trachomatis* infection. Vaccine. 2014;32: 1563–1571.

6. Abdelrahman YM, Belland RJ. The chlamydial developmental cycle. FEMS Microbiol Rev. 2005;29: 949–959.

7. Hybiske K, Stephens RS. Exit strategies of intracellular pathogens. Nat Rev Micro. 2008;6: 99–110.

8. Hybiske K, Stephens RS. Mechanisms of host cell exit by the intracellular bacterium Chlamydia. Proc Natl Acad Sci USA. 2007;104: 11430–11435.

9. Neeper ID, Patton DL, Kuo CC. Cinematographic observations of growth cycles of *Chlamydia trachomatis* in primary cultures of human amniotic cells. Infect Immun. 1990;58: 2042–2047.

10. Chin E, Kirker K, Zuck M, James G, Hybiske K. Actin recruitment to the Chlamydia inclusion is spatiotemporally regulated by a mechanism that requires host and bacterial factors. PLoS ONE. 2012;7: e46949.

11. Lutter EI, Barger AC, Nair V, Hackstadt T. *Chlamydia trachomatis* inclusion membrane protein CT228 recruits elements of the myosin phosphatase pathway to regulate release mechanisms. Cell Rep. 2013;3: 1921–1931.

12. Volceanov L, Herbst K, Biniossek M, Schilling O, Haller D, Nölke T, et al. Septins arrange F-actin-containing fibers on the *Chlamydia trachomatis* inclusion and are required for normal release of the inclusion by extrusion. MBio. 2014;5: e01802–14.

13. Wang Y, Kahane S, Cutcliffe LT, Skilton RJ, Lambden PR, Clarke IN. Development of a transformation system for *Chlamydia trachomatis*: restoration of glycogen biosynthesis by acquisition of a plasmid shuttle vector. PLoS Pathog. 2011;7: e1002258.

14. Tarun AS, Baer K, Dumpit RF, Gray S, Lejarcegui N, Frevert U, et al. Quantitative isolation and in vivo imaging of malaria parasite liver stages. Int J Parasitol. 2006;36: 1283–1293.

15. Sturm A, Amino R, van de Sand C, Regen T, Retzlaff S, Rennenberg A, et al. Manipulation of host hepatocytes by the malaria parasite for delivery into liver sinusoids. Science. 2006;313: 1287–1290.

16. Baer K, Klotz C, Kappe SH, Schnieder T, Frevert U. Release of hepatic Plasmodium yoelii merozoites into the pulmonary microvasculature. PLoS Pathog. 2007;3: e171.

17. Graewe S, Rankin KE, Lehmann C, Deschermeier C, Hecht L, Froehlke U, et al. Hostile takeover by Plasmodium: reorganization of parasite and host cell membranes during liver stage egress. PLoS Pathog. 2011;7: e1002224.

18. Goth SR, Stephens RS. Rapid, transient phosphatidylserine externalization induced in host cells by infection with Chlamydia spp. Infect Immun. 2001;69: 1109–1119.

19. Peterson EM, la Maza de LM. Chlamydia parasitism: ultrastructural characterization of the interaction between the chlamydial cell envelope and the host cell. J Bacteriol. 1988;170: 1389–1392.

20. Doughri AM, Storz J, Altera KP. Mode of entry and release of chlamydiae in infections of intestinal epithelial cells. J Infect Dis. 1972;126: 652–657.

21. Wilson DP, Timms P, McElwain DLS, Bavoil PM. Type III secretion, contact-dependent model for the intracellular development of chlamydia. Bull Math Biol. 2006;68: 161–178.

22. Timms P, Wilson DP, Whittum-Hudson JA, Bavoil PM. Kinematics of intracellular chlamydiae provide evidence for contact-dependent development. J Bacteriol. 2009;191: 5734–5742.

23. Steele LN, Balsara ZR, Starnbach MN. Hematopoietic cells are required to initiate a *Chlamydia trachomatis*-specific CD8+ T cell response. J Immunol. 2004;173: 6327–6337.

24. Matsumoto A. Isolation and electron microscopic observations of intracytoplasmic inclusions containing Chlamydia psittaci. J Bacteriol. 1981;145: 605–612.

25. Criss AK, Seifert HS. A bacterial siren song: intimate interactions between Neisseria and neutrophils. Nat Rev Micro. 2012;10: 178–190.

26. Thwaites GE, Gant V. Are bloodstream leukocytes Trojan Horses for the metastasis of Staphylococcus aureus? Nat Rev Micro. 2011;9: 215–222.

27. Lipkin ES, Moncada JV, Shafer MA, Wilson TE, Schachter J. Comparison of monoclonal antibody staining and culture in diagnosing cervical chlamydial infection. J Clin Microbiol. 1986;23: 114–117.

